# Online narrative guides for illuminating tissue atlas data and digital pathology images

**DOI:** 10.1101/2020.03.27.001834

**Authors:** Rumana Rashid, Yu-An Chen, John Hoffer, Jeremy L. Muhlich, Jia-Ren Lin, Robert Krueger, Hanspeter Pfister, Richard Mitchell, Sandro Santagata, Peter K. Sorger

**Affiliations:** Laboratory of Systems Pharmacology, Harvard Medical School, Boston, MA; Ludwig Center for Cancer Research at Harvard, Harvard Medical School, Boston, MA; Department of Pathology, Brigham and Women’s Hospital, Harvard Medical School, Boston, MA; School of Engineering and Applied Sciences, Harvard University, Cambridge, MA; Department of Systems Biology, Harvard Medical School, Boston, MA

## Abstract

The recent development of highly multiplexed tissue imaging promises to substantially accelerate research into basic biology and human disease. Concurrently, histopathology in a clinical setting is undergoing a rapid transition to digital methods. Online tissue atlases involving highly multiplexed images of research and clinical specimens will soon join genomics as a systematic source of information on the molecular basis of disease and therapeutic response. However, even with recent advances in machine learning, experience with anatomic pathology shows that there is no immediate substitute for expert visual review, annotation, and description of tissue images. In this perspective we review the ecosystem of software available for analysis of tissue images and identify a need for interactive guides or “digital docents” that allow experts to help make complex images intelligible. We illustrate this idea using *Minerva* software and discuss how interactive image guides are being integrated into multi-omic browsers for effective dissemination of atlas data.

## MAIN TEXT

Light microscopy, most commonly transmission microscopy of specimens stained with colorimetric dyes, is a traditional, powerful, and widely used method for diagnostic pathology and for studying tissues in research. The relatively recent introduction of highly multiplexed tissue imaging_1–4_ now makes it possible to conduct deep molecular profiling of tissues and tumors at subcellular resolution while preserving native tissue architecture^5^. Multiplexed, high resolution imaging greatly facilitates the study of cell states, cell-cell interactions, and tissue architecture in normal and disease conditions. A common application of tissue imaging in oncology, for example, is identifying and enumerating immune cell types and mapping their locations relative to tumor and stromal cells^5^. Such spatially-resolved data is pertinent to understanding the mechanisms of action of immunotherapies (e.g., anti-PD1 or PDL1 immune checkpoint inhibitors)^6^ that function by blocking juxtracrine signaling between immune and tumor cells. Tissue images are complex however: biologically relevant structures range in size by over five orders of magnitude from subcellular vesicles and nuclear granules at micron scales to configurations of cells in blood and lymphatic vessels at millimeter scales to interactions among endothelia, epithelia, muscle and other cell types to form functioning tissues at centimeter scales.

A variety of multiplexed tissue imaging methods has been described over the last few years. These include Imaging Mass Cytometry (IMC)^1^ and Multiplexed Ion Beam Imaging (MIBI)^2^, which detect antigens using metal isotope-labeled antibodies, tissue ablation, and atomic mass spectrometry. In contrast, methods such as MxIF^3^, CODEX^7^, t-CyCIF^4^, multiplexed IHC^8,9^, and immuno-SABER^10^, use fluorescently-labelled (or enzyme-linked) antibodies followed by imaging on microscopes. These methods differ in the number of antigens they can routinely detect on a single tissue section (currently ~12 in the case of multiplexed IHC to ~40-60 in the case of IMC or t-CyCIF). Some methods are restricted to selected fields of view (e.g., ~500 µm square for MIBI and IMC) whereas others can perform whole slide imaging (WSI) on areas ~400-1,000 times larger (e.g. -CyCIF or CODEX). Most multiplexed tissue imaging methods are in active development and their strengths and limitations with respect to speed, sensitivity, resolution, etc. remain to be determined. However, they all generate multiplexed 2D images of cells and supporting tissue structures *in situ*. When data are collected using high resolution microscopes it is also possible to generate 3D images by optical sectioning and tissue clearing methods^11^.

### Tissue and tumor atlases

International projects are currently underway to create publicly accessible atlases of normal human tissues and tumors. These include the Human Cell Atlas^12^, the Human BioMolecular Atlas Program (HuBMAP)^13^ and the Human Tumor Atlas Network (HTAN)^14^. For example, HTAN is envisioned to be a spatially resolved counterpart of the well-established Cancer Genome Atlas (TCGA)^15^ and Encyclopedia of DNA Elements (ENCODE)^16^ (**Figure 1**). HTAN atlases aim to combine the genetic and molecular precision of dissociative single-cell methods such as single-cell RNA sequencing with morphological and spatial information obtained from tissue imaging and spatial transcriptomics^13,14^. At their inception, the imaging components of these atlases are likely to contain data acquired from one or a few individuals but they will ultimately merge multiple specimens into common reference systems^13^. Conceptually, integration across samples and data types is easiest to imagine at the level of *derived features*, such as a census of cell types and positions (from imaging data) or transcript levels (from scRNA-Seq). Adding meso-scale information from images such as the number and arrangement of supporting stroma, membranes, blood and lymphatic vessels, etc. is a greater challenge. It is not yet known how best to capture such spatial information computationally; insights gleaned from centuries of study of tissue by anatomic pathologists therefore remain essential. Studies are underway to better understand how pathologists make diagnoses from tissue specimens^17^ and to quantify connections between features computed from cellular neighborhoods and clinical outcome^18^. However, human inspection of tissue images will almost certainly remain essential for relating morphology to physiology and pathophysiology and also for assessing the quality of image processing algorithms, training classifiers, etc. We therefore require software that allows atlas users to benefit from the expertise of pathologists who currently work almost entirely with physical specimens on glass slides (typically, with the goal of making real-time diagnostic decisions). We envision software interfaces that serve as digital docents that guide users through an expert description of a specimen while also facilitating access to quantitative information derived from computational analysis.

**Figure 1.**
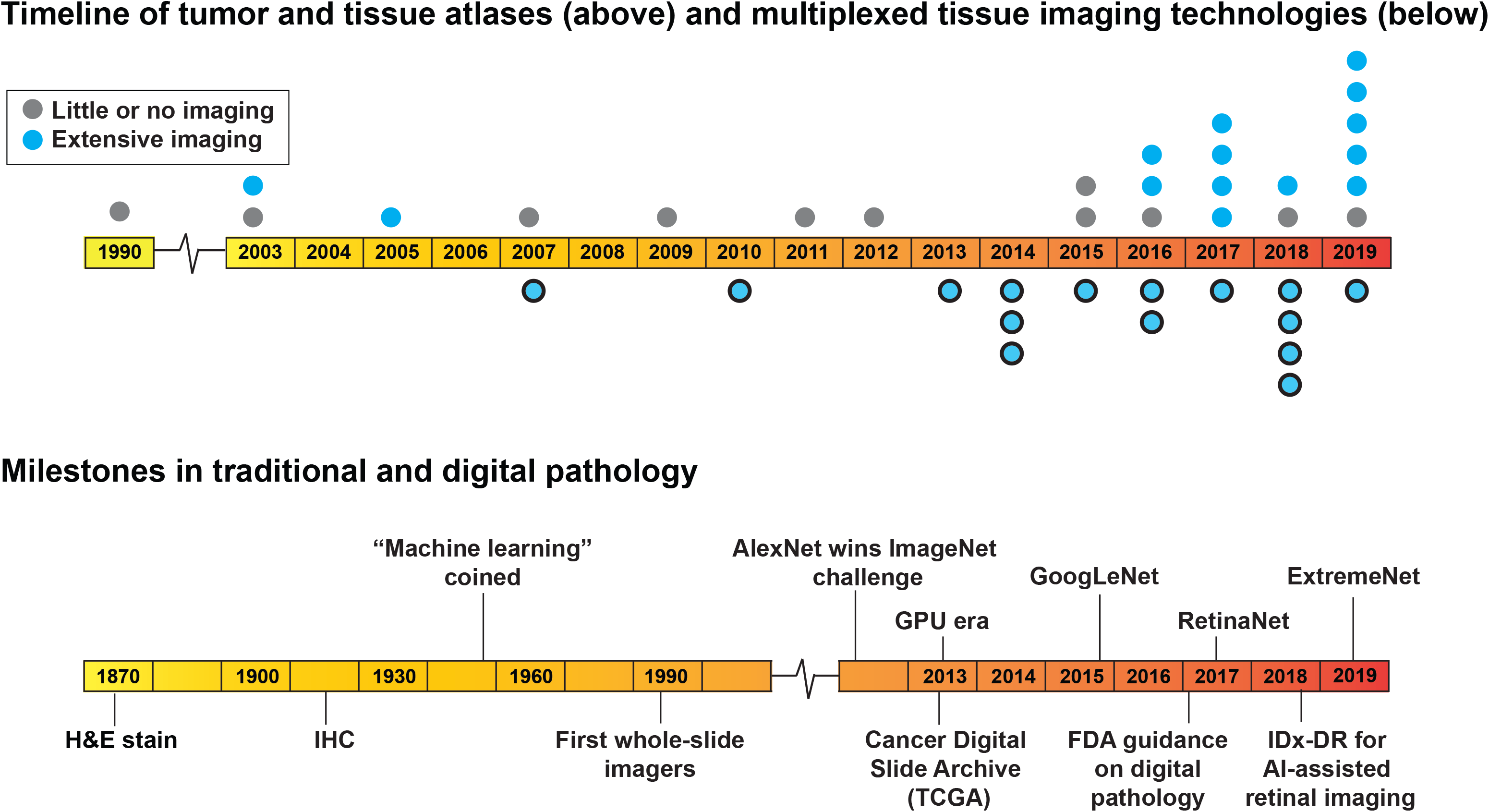
Timeline of milestones in development of human atlases, histopathology, and multiplexed tissue imaging technologies. The upper timeline depicts the establishment of publicly available atlases of human tissues, tumors, and cell types (gray denotes genome-focused atlases and blue denotes atlases with a substantial imaging component). Each dot represents the year that one new large-scale atlas was established. The number of atlases is increasing, as is the emphasis on imaging data. The lower timeline depicts the evolution of methods used for histology and anatomical pathology, with an accelerating trend towards digital methods occurring in parallel with growth of multiplexed imaging and atlases. AI: Artificial intelligences; IHC: immunohistochemistry; FDA: Food and Drug Administration; GPU: graphics processing unit; TCGA: the Cancer Genome Atlas; IDx-DR: an AI-based diagnostic for diabetic retinopathy from www.eyediagnosis.net.

### Tissue imaging in a clinical setting

The introduction of multiplexed tissue imaging in a research setting is occurring concurrently with an accelerating transition to digital technologies^19^ in clinical histopathology (**Figure 1**). While genetics is pertinent to disease diagnosis, particularly in cancer and inherited diseases, histology and cytology remain the central pillars of routine clinical work and are one of the primary diagnostic modalities for most diseases. In current practice, tissue samples recovered by biopsy or surgical resection are formaldehyde-fixed and paraffin embedded (FFPE), sliced into 5µm sections, and stained with hematoxylin and eosin (H&E). Samples for immediate study, when a patient is still undergoing surgery for example, are frozen, sectioned, and stained (these are often called OCT samples based on the “optimal cutting temperature” medium in which they are embedded). Liquid hematologic samples are spread on a slide to create blood smears, which are also stained with colorimetric dyes (Romanowsky– Giemsa staining). H&E staining imparts a characteristic pink to blue color on cells and other structures and pathologists review these specimens using simple bright-field microscopes; other stains are analyzed in a similar way. Some clinical samples are also subjected to immunohistochemistry (IHC) to obtain information on the expression of one or a small number of protein biomarkers per slide^20^. While cost-effective and widely used, many histopathology methods were developed over a century ago, and IHC is itself 75 years old^21^. Moreover, diagnoses based on these techniques generally do not capture the depth of molecular information needed to optimally select targeted therapies. The latter remains the purview of mRNA and genome sequencing which, in a clinical setting, often involves exome sequencing of selected genes.

Recently, pathologists have started to leverage machine learning to assist with pattern recognition from histologic data and potentially extract deeper diagnostic insight. Digital analysis of histological specimens first became possible with the introduction of bright-field WSI instruments twenty years ago^22,23^, but it was not until 2016 that the FDA released guidance on the technical requirements for use of digital imaging in diagnosis^24^ (**Figure 1**). Digital pathology instruments, software, and startups have proliferated over the past few years fueled in large part by the development of machine learning algorithms capable of assisting in the interpretation of H&E stained slides^25^, which histopathology services must process in very high volume (often >1 million slides per year in a single hospital). Machine learning on images has proven successful in several areas of medicine^26,27^ and promises to assist practitioners by increasing the efficiency and reproducibility of pathologic diagnosis^25^. The pathology departments at several NCI comprehensive cancer centers have recently introduced multiplexed image-based immunoprofiling services to identify patients most likely to benefit from immune checkpoint inhibitors. Nonetheless, in most hospitals, the vast majority of diagnostic pathology still involves visual inspection of physical specimens; only a minority of slides are scanned and digitized for concurrent or subsequent review on a computer screen. This is widely anticipated to change over the next decade and several European countries are implementing national digital pathology programs. As clinical pathology incorporates new measurement modalities and becomes increasingly digital, it is highly desirable that software and standards developed for clinical and research purposes be compatible and interoperable; this is likely to be particularly important in the conduct of clinical trials. From a technical perspective, standards such OME-TIFF developed for pre-clinical research work well with multiplexed images from clinical services. However, software for clinical use requires security, workflow and billing features that are not components of research systems.

### Accessing and sharing imaging data in tissue atlases

Algorithms, software, and standards for high-dimensional image data^28^ remain under-developed relative to tools for almost all types of genomic information. Moreover, with sequencing data, the information present in primary data files (e.g., FASTQ files) are fully retained (or enhanced) when reads are aligned or count tables generated; it is rarely necessary to re-access the primary data. In contrast, methods to extract features from tissue images are immature, and visual inspection by knowledgeable viewers as well as development and testing of new algorithms requires access to image data at native resolution. Unfortunately, the software and computational resources needed to make cell and tissue images broadly available have been slow to develop.

As far back as 2008, the *Journal of Cell Biology* worked with the founders of the Open Microscopy Environment (OME)^29^ to deploy a JCB DataViewer^30^ that provided direct access to primary, high-resolution microscopy data (much of it from tissue culture cells and model organisms). Economic pressures ended this ambitious effort^31^, emphasizing that funds have long been available to purchase expensive microscopes, but not to distribute the resulting data. Currently, most H&E, IHC, and multiplexed tissue images are shared only as figure panels in manuscripts, a form that typically provides access to a few selected fields of view at a single resolution. In the best case these data might be available via Figshare^32^ although the European Bioinformatics Institute Image Data Resource (IDR) is a notable exception; IDR uses the OME-compatible OMERO server to provide full-resolution access to selected microscopy data^33^. The importance of sharing image data is particularly pronounced in the case of research biopsies, including those being used to assemble tumor atlases. These biopsies are obtained to advance scientific knowledge rather than inform the treatment of individual patients and there is an ethical obligation for the resulting data (appropriately anonymized) to be made available in an open and useful form that accelerates scientific discovery^34^. This principle is widely recognized in genomics^35,36^ but has only recently been addressed in the area of tissue imaging^37^. More generally, digital pathology and tissue imaging are disciplines in which the goal of making research Findable, Accessible, Interoperable, and Reusable (FAIR)^38^ is highly relevant, but the necessary computational infrastructure is deficient.

### Software for image analysis, management, and interpretation

To a first approximation, the wide variety of academic and commercial microscopy software currently available has been implemented either as a desktop system focused on data analysis or, like OMERO, as a client-server relational database management system (RDBMS) system focused on image management (**Figure 2a**; see **Box 1** and **Box 2** for details). Desktop software is particularly good for interactive image analysis because it exploits graphics cards for rapid image rendering and high-bandwidth connections to local data for computation. RDBMS systems are ideal for data management because they enable relational queries, support multiple simultaneous users, ensure data integrity, and effectively manage access to large-scale local and cloud-based compute resources (a more detailed comparison of available software is provided in **Figure 2b**).

**Figure 2.**
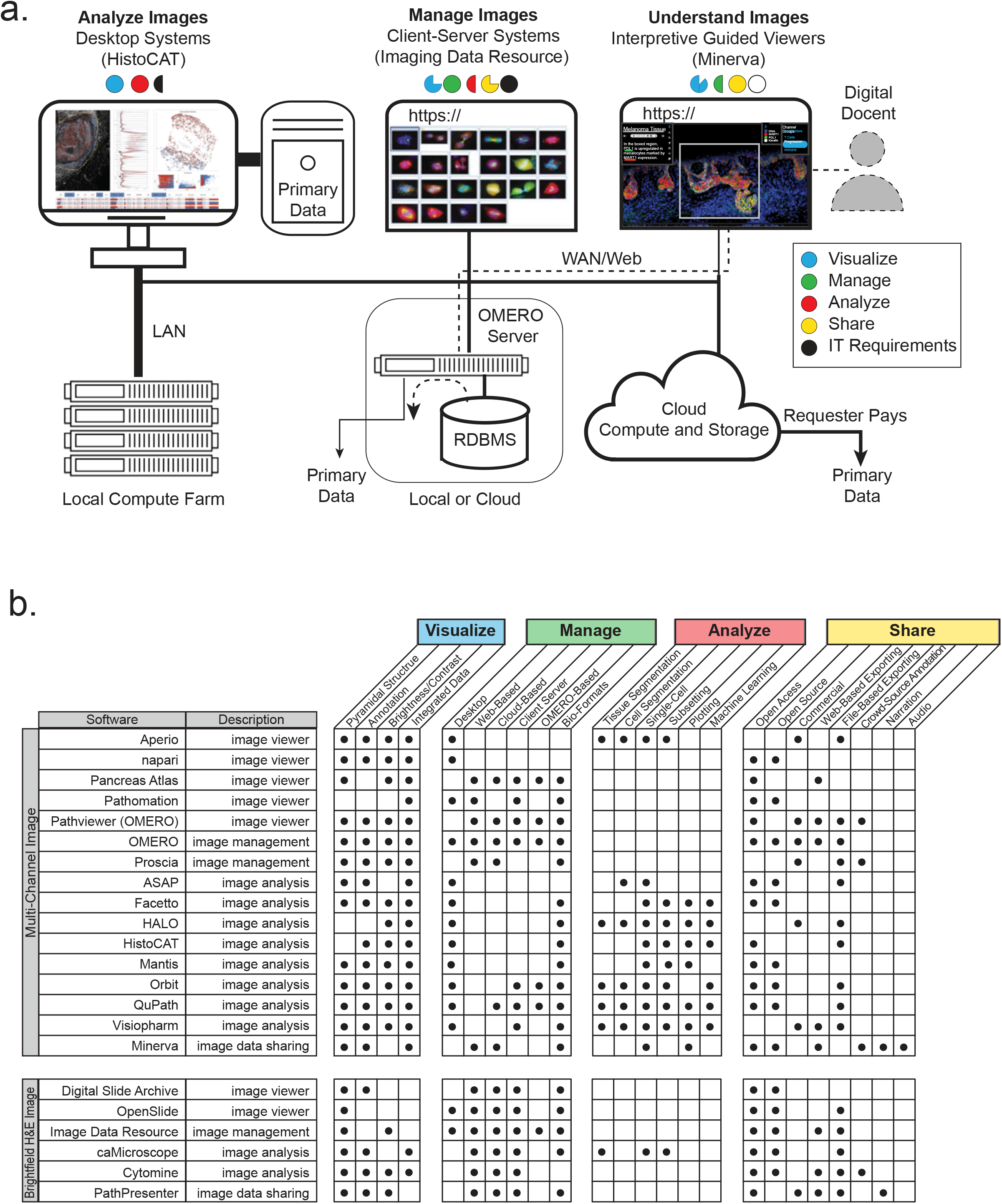
Software used to visualize, analyze, manage, and share tissue images. A variety of software is available for processing tissue image data, each with strengths and weaknesses. We expect them to co-evolve as part of an ecosystem held together by common data standards and interoperable application programming interfaces (APIs). **(a)** Desktop applications such as *histoCat* provide sophisticated tools for interactive quantification and analysis of image data and effectively exploit embedded image rendering capabilities. Many desktop applications can be run in a “headless” configuration on compute farms to accelerate analysis of large images. They can also fetch images from *OMERO*. Three-tier client server systems such as *OMERO* involve a relational database management system (RDMS), an image server, and one or more browser-based and desktop clients; such systems provide a full-featured approach to organizing, visualizing, and sharing large numbers of images but they typically support only limited data processing. The *Minerva* application described in this perspective is optimized for data sharing and interpretative image viewing: within *Minerva* detailed narration of images and derived data is possible. Pre-rendering of stories with multiple waypoints in an image mimics the insight provided by an expert “docent”. Public access to large-scale primary data is typically limited with all of these systems because of the costs of data transfer. In some OMERO configurations (e.g. the Image Data Resource) primary data can be downloaded from the same file system. Another approach to primary data access being used by HTAN involves “requester pays” cloud-based storage (e.g., an Amazon Web Service S3 bucket). *Minerva Author* is currently being extended so it can access data stored in OMERO. **(b)** The range of commercial and academic software suitable for viewing image data with key features described. While each tool has its strengths and weaknesses, none of these tools comprehensively satisfies all of functions needed for image analysis. We expect data generators and consumers to rely on a suite of interoperable desktop and server or cloud-based software.

However, as the first high-plex, whole-slide tissue images have become available it has become clear that a new type of software will be required to guide users through the extraordinary complexity of images that encompass multiple square centimeters of tissue, 10^5^-10^7^ cells, and upwards of 100 channels. We envision a key role for “interpretative guides” (digital docents) that help walk users through a series of human and machine-generated annotations about an image in much the same way that the results section of a paper guides users through a multi-panel figure. Genomic science faced an analogous need for efficient and intuitive visualization tools a decade ago and this lead to the development of the highly influential Integrative Genomics Viewer^39^ and its many derivatives.

Interactive guides of images have proven highly successful in other fields, as exemplified by *Project Mirador* (https://projectmirador.org/). *Project Mirador* focuses on the development of web-based interpretative tours of cultural resources such as art museums, illuminated manuscripts, and culturally or historically significant cities. In such online tours, a series of waypoints and accompanying text direct users to areas of interest while also allowing free exploration and a return to the narrative.

This mimics the functionality of museum guides and docents. Interactive narration (also known as digital storytelling or visual storytelling) has also demonstrated strengths as a pedagogical tool that enhances comprehension^40^ and memory formation^41^. Multiple studies have identified benefits associated with receiving complex information in a narrative manner, and digital storytelling has been applied to several areas of medicine and research including oncology^42^, mental health^43^, health equity^44^, and science communication^45,46^.

How might these lessons be applied to tissue images? At the very least they teach us that it is insufficient to simply make gigapixel-sized images available for download and analysis on desktop software. Instead, we require an easy way to guide users through the salient features of an image with associated annotation and commentary. Watching a pathologist describe a specimen to a colleague provides additional insight. She or he typically uses a multi-head microscope to pan across an image and switch between high and low power fields (magnifications), thereby studying cells in detail while also placing them in the context of the overall tissue. In this process, key features are often highlighted using an LED pointer. With multiplex images, it is also necessary to toggle channels on and off so that the contribution of specific antibodies to the final image can be ascertained.

### Software-based interpretive guides for sharing tissue images

With these requirements in mind, we asked how interpretive guides for tissue images might be implemented. One obvious possibility is as an OMERO client. OMERO is the most widely deployed open source image informatics system and it is compatible with a range of software clients. However, an OMERO client requires access to an OMERO database and, as images get larger, the server becomes substantially loaded, limiting the number of concurrent sessions. We therefore settled on a database-independent viewer based on the open source OpenSeadragon^47^ platform used by *Project Mirador* and other software tools. OpenSeadragon makes it easy to zoom and pan across images in manner similar to Google Maps^48^. The resulting *Minerva* software^49^, is a single-page web application that uses client-side JavaScript and is easily deployable on standard commercial clouds (e.g., Amazon Web Services - AWS; **Figure 3**) or on local computing servers that support Jekyll^50^. OpenSeadragon has been used previously for displaying H&E images^51^, but in *Minerva* it is paired with narrative features, interactive views of derived single-cell features within the image space, a lightweight implementation, and the ability to accommodate both bright-field and multiplexed immunofluorescence images.

**Figure 3.**
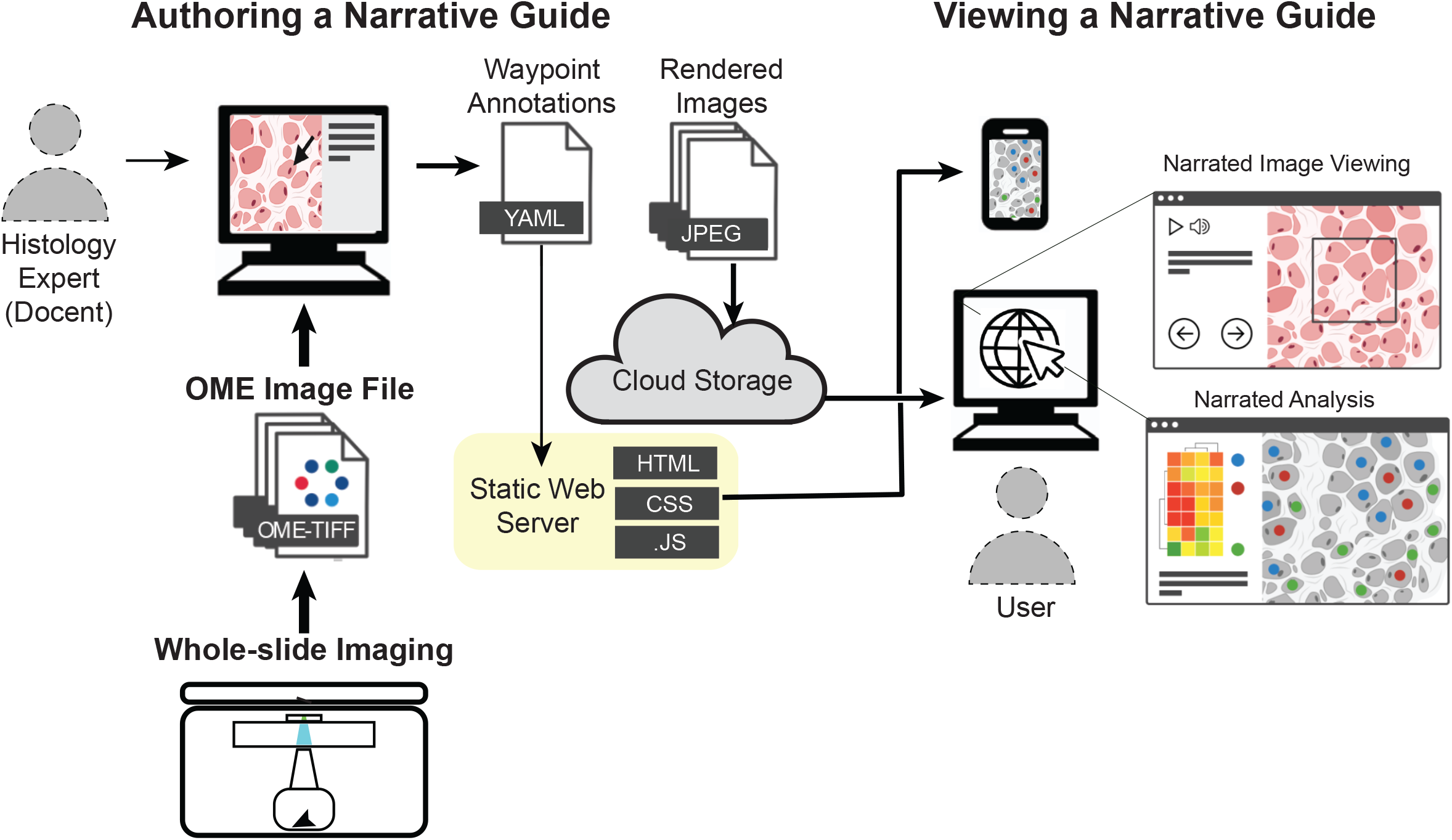
A system for generating and viewing on-line narrative guides to complex tissue images. The architecture illustrated here is based on the OpenSeadragon viewer *Minerva Story* and the narration tool *Minerva Author*; the thickness of the arrows indicates the amount of data transferred. Whole slide images including bright field and multi-channel microscopy images in the standard OME-TIFF format are imported into *Minerva Author* by a user with expertise in tissue biology and image interpretation (the docent); in many cases this individual is a pathologist or histologist. The docent uses tools in *Minerva Author* to pan across the image and then set channel rendering values (background levels and intensity scale), specify waypoints, and add text and graphical annotations. *Minerva Author* then renders image pyramids for all channel groupings as JPEG files and generates a YAML configuration file that specifies waypoints and associated information. The rendered images are stored on a cloud host (e.g., Amazon Web Services S3) and accessed via a static web server supporting Jekyll, on GitHub for example. A user simply opens a web browser, clicks on a link, and *Minerva Story* launches the necessary client-side JavaScript (.JS), making it possible to follow the story and also freely explore the image through pan, zoom, and channel selection actions. Because interactivity is handled on the client, no special backend software or server is needed. Images rendered by *Minerva Story* are compatible with multiple devices, including cell phones.

Minerva is OME- and BioFormats-compatible and therefore usable with images from virtually any existing microscope or slide scanner. There is no practical limit to the number of users who can concurrently access narrative guides developed in *Minerva* and no requirement for specialized servers or relational database, keeping complexity and costs low. Anyone familiar with GitHub and AWS or similar cloud services can deploy a *Minerva* story in a few minutes, and new stories can then be generated by individuals with little expertise in software or computational biology. Developers can also build on the *Minerva* framework to develop narration tools for other applications such as protein structures or complex high-dimensional datasets. Viewers such as *Minerva* are not intended to be all-in-one solutions to the many computational challenges associated with processing and analyzing tissue data. Instead, they are specialized browsers that perform one task well: in the case of *Minerva*, by providing an intuitive, interpretative approach to image data. This liberates images from static and ‘postage-stamp’ representations in journals. Genome browsers are similar: they do not perform alignment and data analysis, but they make it possible to interact with processed sequences effectively.

Within a *Minerva* window on a standard Web browser, a narration panel directs a user’s attention to particular regions of an image and specific channel groupings, accompanied by text description (which *Minerva* can read aloud) as well as image annotations involving overlaid rectangles, polygons, arrows, text, etc. (**Figure 4**). Each image can be associated with more than one narrated story and with different ways of viewing the same type of data. A fundamental aspect of narrative guides is that individuals with expertise in a particular disease or tissue, such as a pathologist, create narratives used by others to assist in understanding the morphology of a specimen. Creating narrations requires an authoring tool such as *Minerva Author*. *Minerva Author* is a desktop application (in JavaScript React with a Python Flask backend) that converts images in OME-TIFF^52^ format into pyramidal form and assists with the addition of waypoints and text annotations. *Minerva Author* supports RGB images (brightfield, H&E, immunohistochemistry, etc.) as well as multi-channel images (immunofluorescence, CODEX, CyCIF, etc.). After specifying rendering settings and writing the waypoints in *Minerva Author*, a user receives a configuration file and image pyramids to deploy to AWS S3 or another Web-based storage location (**Figure 3**). Stories can be as simple as a single panel with a short introduction or a multi panel narration enriched with a series of views with detailed descriptions, changes in zoom level, and associated data analysis. We find that it takes users a few hours to learn the software and then 30 minutes to a few hours to create a story, about the same time required to create a good static figure panel for a journal and much less time than data collection, image registration, segmentation, and data analysis.

**Figure 4.**
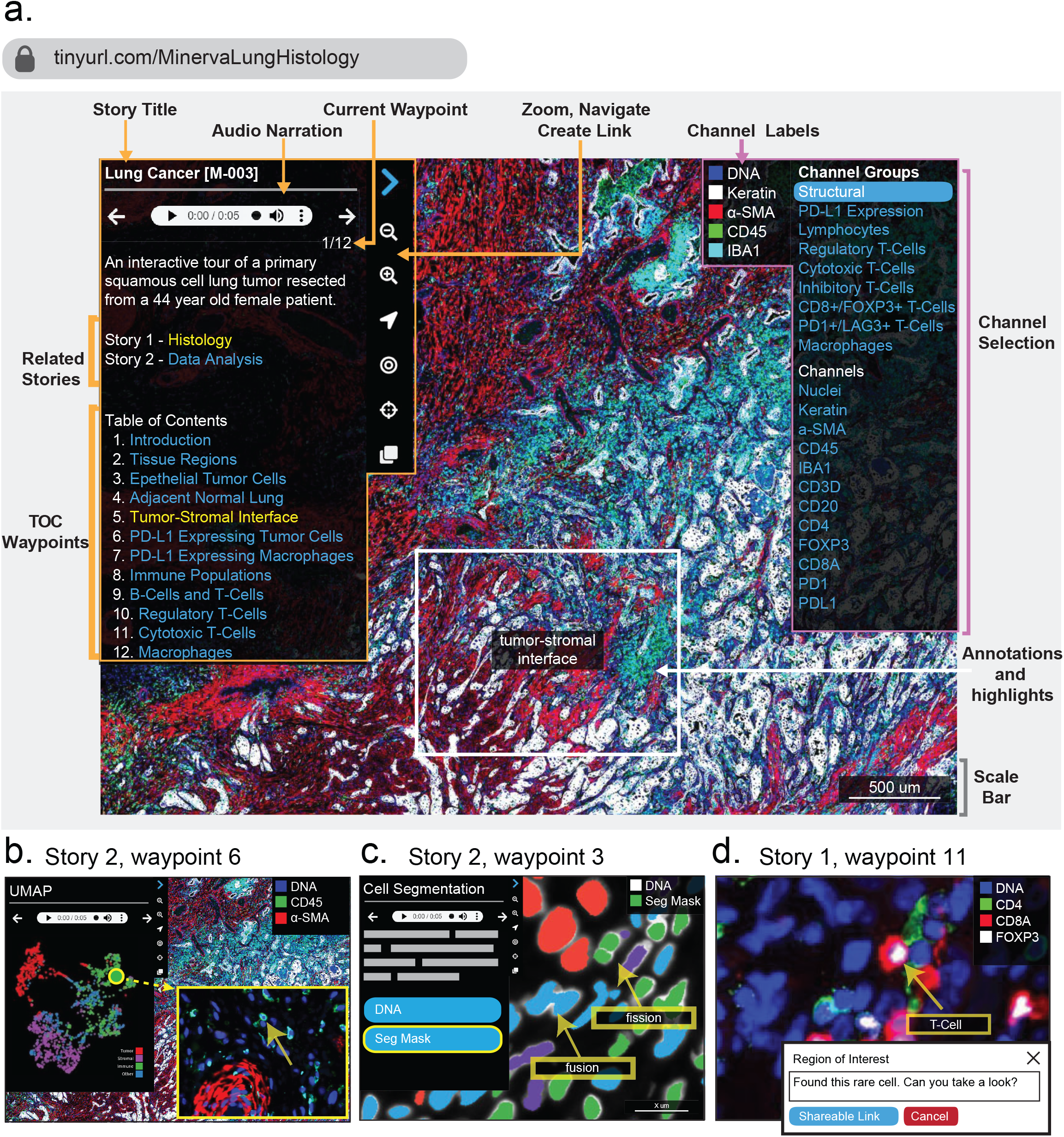
Minerva Story. The *Minerva Story* user interface showing key features. Note that this is a synthetic screenshot, and size and rendering of the text has been altered from the original to make the interface easier to understand in a figure panel. Two *Minerva* stories are available for this multiplexed image of a lung adenocarcinoma specimen. Story one, focusing on histology and immune populations, can be accessed at https://tinyurl.com/MinervaLungHistology and story two, focusing on data generation and analysis, at https://tinyurl.com/MinervaLungData. **(a)** The home screen for *Minerva* story one. A narration panel on the left-hand side shows the title and narrative text that can also be read aloud using the audio panel. The left panel also has links to related stories for the specimen and a table of contents listing each waypoint in the story (outlined in orange). Users can click the navigation arrows to step through each waypoint or skip to a waypoint of interest using the table of contents. On the right side is channel selection panel that users can use to change which channels are rendered (outlined in pink). At any point, users can depart from a story and freely pan and zoom in or out of the image using the mouse or the magnification icons in the navigation panel. Arrowheads, concentric circles, and other graphical elements can be used to annotate images and generate web URLs specific to the current rendering. The viewer also contains a scale bar that grows and shrinks as a user navigates across zoom levels (grey). Note that this synthetic screenshot includes only a subset of the waypoints and channel groups. Extensive information on these features can be found at https://github.com/labsyspharm/minerva-story/wiki. **(b)** Waypoint 6 from story two displays an interactive plot of a UMAP (Uniform Manifold Approximation and Projection for Dimension Reduction) performed on a random sample of 2,000 cells from the tissue. Users can click on any data point in the plot (each data point represents a single cell) and the browser will zoom to the position of that cell in the image and place an arrow. **(c)** Waypoint 3 from story two shows each cell in the tissue overlaid with a segmentation mask in which the color denotes the cell type as determined by quantitative k-means clustering. This makes it possible for users to assess the accuracy and effects of segmentation on downstream cell type calling analysis. The reader can toggle the mask with immunofluorescence channels to view the quantitative and image data in the same view. The waypoint annotates two types of common segmentation errors. **(d)** Demonstration of on-the-fly annotation tool; users can add annotations and enter text and then hit the blue button to generate a URL that allows anyone to render the same image and location with pan, zoom, marker group, and annotations preserved.

### Using interactive guides to explore a human tumor

As a use case for interactive tissue guides, we used *Minerva Story* to mark up a large human lung adenocarcinoma specimen (tinyurl.com/MinervaLungHistology). This primary tumor measured ~ 5 mm x 3.5 mm and was imaged at subcellular resolution using 44-plex t-CyCIF^53^; multiple fields were then stitched into a single image as described in detail in a recent publication^54^ (the image is referred to as “LUNG-3-PR”). There is no single best way to analyze an image containing several hundred thousand cells and we therefore created two guides focused on (i) histologic review of regions of interest and specific immune and tumor cell types and (ii) presentation of quantitative data analysis in the context of the original image (**Figure 4a**). The name of the samples, links to related stories, a table of contents, and navigation tools are found in a panel to the left of the *Minerva Story* window (outlined in orange in **Figure 4a**). Another navigation panel, focused on channel selection, is found on the right of the window (outlined in pink); channels and channels groups, as well as the cell types they define, can be pre-specified. We typically link each protein (antigen) name to an explanatory source of information such as the GeneCards^55^ database, but a more customized annotation of markers specific to a particular tissue would be preferable and is under development.

Stories progress from one waypoint to the next (the analogy is to numbers used by museum audio guides) and each waypoint can involve a different field of view, magnification, and set of channels, as well as arrows and text describing specific features of interest (marked in gray). At any point, users can diverge from a story by panning and zooming around the image or selecting different channels, and then return to the narrative by clicking on the appropriate waypoint in the table of contents. The field shown in **Figure 4a**, (waypoint one) shows pan-cytokeratin positive tumor cells growing in chords and clusters at the tumor-stromal interface. This region of the tumor is characterized by an inflammatory microenvironment as evidenced by the presence of a variety of lymphocyte and macrophage populations distinguishable by expression of cell surface markers. Using the panel on the right, users can toggle these markers on and off to explore the images and data themselves and evaluate the accuracy of classification. The remainder of the story explores the expression of PD-L1 (Programmed death-ligand 1), an immune checkpoint protein and drug target, and localization of lymphocyte and macrophage populations.

Narrative guides are also useful in showing the results of quantitative data analysis in the context of the original image (this constitutes the second story about the lung adenocarcinoma). Data analysis of high-plex tissue images typically involves measuring the expression of multiple cell type markers (e.g., immune linear markers) in single cells and using this data to identify cell types. Analysis of cell state, morphology, and neighborhood relationships are also common. Within *Minerva Story* it is possible to link representations of quantitative data directly to the image space. For example, when data are captured in a two-dimensional plot, such as a UMAP (**Figure 4b**), clicking on a data point takes the user directly to the corresponding position of the cell in the image (which is denoted with an arrow). This is a standard feature in desktop software such as *HistoCat* and *Facetto* and greatly enhances a user’s understanding of the relationship between images and image-derived features. Display of segmentation masks as overlays is similarly useful for troubleshooting and assessing data quality; in **Figure 4c**, unwanted fusion and fission events are highlighted by arrows, both of which result from errors in segmentation. Additionally, users can interactively highlight areas of interest, add notes, and generate sharable links that allow others to navigate to the same position in the image and view any added annotations and text (**Figure 4d;** in this case a high magnification view is shown with a T-cell labelled).

### Narrative guides as a medical education tool

Teaching is another application for narrative tissue guides. Histology is challenging to teach in an undergraduate setting^56^ and, in the case of medical students and residents, changes in curriculum have resulted in much less time in front of a microscope. On-line collections of tissue images are a frequent substitute. However, studies have shown that pairing classroom instruction with dynamic viewing and flexible interaction with image data are essential for learning^57^. We have therefore created a narrative guide to H&E images of specimens from the heart of a patient who experienced multiple episodes of myocardial infarction prior to death (**Figure 5**). An introductory panel depicts the overall structure of the heart and the positions from which various specimens were resected. These images reveal the histologic hallmarks of ischemic heart disease such as severe coronary artery atherosclerosis, plaque rupture, stunned myocardium, reperfusion injury, and the early, intermediate, and late features of myocardial tissue infarction. The interactive narration of this common clinical syndrome provides a context for developing a more nuanced understanding of cardiac pathophysiology than looking at snippets in a textbook or poorly annotated on-line images.

**Figure 5.**
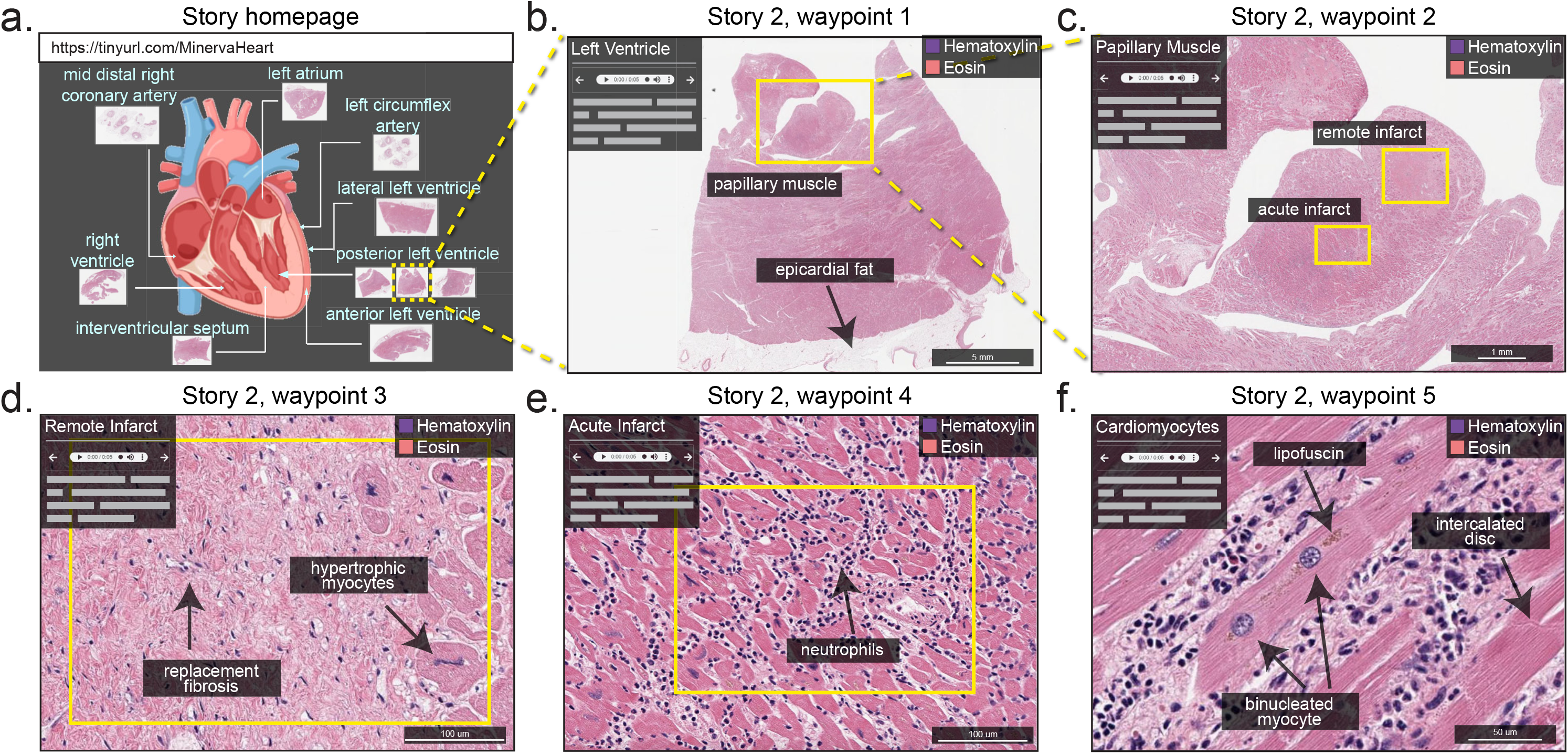
An application of *Minerva Story* in medical education. *Minerva Story* can be used for multiple applications, including guided viewing of conventional H&E stained sections. In this example available at https://tinyurl.com/MinervaHeart, a *Minerva* story has been created to guide students through tissue specimens from different anatomical regions of a heart from a patient who suffered multiple myocardial infarcts. **(a)** An anatomic schematic of the heart indicates the regions from which tissue specimens in the *Minerva* story were collected. **(b)** Waypoint 1 shows a specimen sliced with a posterior view of the left ventricle with box, arrow, and text annotations of a few histologic features. **(c)** Waypoint 2 of the story in (b) showing a zoomed-in view of a papillary muscle characteristic of the left ventricle with annotations indicating regions of tissue showing acute (~12 days old) and remote (>6 weeks to years old) myocardial infarct. **(d)** Waypoint 3 and **(e)** Waypoint 4 guide the user to the remote infarct and acute infarct regions of the papillary muscle describing histologic hallmarks to distinguish the two types of infarcts. **(f)** Waypoint 5 depicts microscopic cellular structures of cardiomyocytes at high resolution in a region of the tissue with a late acute infarct. This *Minerva* story is available at https://tinyurl.com/MinervaHeart.

As the practice of digital pathology continues to grow, we anticipate that uptake of software such as *Minerva Story* in the medical education community could help with the creation of interactive textbooks. With relatively little effort, existing text can be hyperlinked to stories and waypoints and students can use image annotation features to take notes and ask questions. Audio based narration is also advantageous in this context as it allows students to receive lecture content while simultaneously concentrating on an image.

### Outlook

Multiplex tissue imaging methods generate images that are data-rich but the potential for this data to inform basic, translation, and clinical research remains largely untapped, in large part because the complexity and size of such images makes them difficult to process and share. Demand for data access will grow as atlases come on-line and the first papers using atlas data are published. Software is therefore required to capture and disseminate images in an interactive form, similar to the time-honored practice of a pathologist sitting down with a colleague or student to describe a specimen, while also enabling free data exploration. With cloud-based storage of pre-rendered narrative guides, there is no real limit to the number of people who can view an image and costs for data providers are low. These possibilities are illustrated by *Minerva* and the OpenSeadragon platform on which it is based. *Minerva* is also capable of displaying images on a cell phone, which is useful in both educational and clinical environments.

With respect to data analysis, a long-term goal in computational biology is to have one copy of the data in the cloud and then move computational algorithms to the data. At present however, local analysis of images remains the dominant approach, particularly for interactive tasks such as optimizing segmentation, training machine learning algorithms, performing and validating clustering, etc. This requires local access to full-resolution primary data. With commercial cloud services, data egress (download) is the primary expense, and can be substantial for multi-terabyte datasets. A likely solution is the use of “requester pays” buckets available on Amazon, Google, and other cloud services. These allow a data generator to make even large datasets publicly available by requiring the requester to cover the cost of data transfer. In the short term, we envision an ecosystem in which web clients for databases such OMERO provide some interactive analysis of primary data, with private datasets allowing for more functionality than public datasets because demand on the server can be better managed. This ecosystem would include lightweight viewers such as *Minerva* that provide access to published and processed data – and make data intelligible to non-experts – and requester pays cloud buckets allowing computational biologists to access primary data and perform novel analysis locally.

In the longer term, these functions are likely to merge, with OMERO providing sophisticated cloud-based processing of data and *Minerva* serving as a means to access data either in OMERO or a static file system virtually cost-free. One obvious extension of *Minerva* is to add tools that facilitate supervised machine learning; in this application, a *Minerva* variant provides an efficient way of adding the labels to data needed to train classifiers and neural nets. The results of machine classification can also be checked in *Minerva*, whose lightweight implementation facilitates crowdsourcing. With these possibilities in mind a key question is whether tools such as *Minerva* should expand to include sophisticated image processing functions. We think not; instead we believe the narrative guides built using *Minerva* (or other tools to be developed) should be optimized for image review, publication, and description. Analysis will continue to be performed using other (interoperable) software tools. These can be joined together into efficient workflows as needed using software containers (e.g. Docker)^58^ and pipeline frameworks such as Nextflow^59^. As described in Box 2, this approach is not perfect^60^, but it cannot reasonably be replaced by all-in-one commercial or academic software.

Recent interest in single-cell tissue biology derives not only from advances in microscopy but also from the widespread adoption of single cell sequencing^61,62^. Comprehensive characterization of normal and diseased tissues will almost certainly involve the integration of data from multiple analytical modalities, including imaging, spatial transcript profiling^63,64^, mass spectrometry imaging of metabolites and drugs^65,66^, and computational registration of dissociated scRNA-seq^67^ with spatial features. *Minerva* cannot handle all of these tasks but it can be readily combined with other tools to create the multi-omic viewers needed for tissue atlases. An immediate goal is adding narrative tissue guides to widely used genomics platforms such as cBioPortal for Cancer Genomics^68^ to create environments in which genomic and multiplexed tissue histopathology can be viewed simultaneously. Better visualization will also help with the more conceptually challenging task of integrating spatio-molecular features in multiplex images with gene expression and mutational data. With respect to clinical applications, the biomedical community needs to ensure that digital pathology systems do not become locked behind proprietary data formats based on non-interoperable software. OME and BioFormats for microscopy and DICOM for radiology^69^ demonstrate that it is possible for academic developers, commercial instrument manufactures and software vendors to work together for mutual benefit. Easy to use software such as *Minerva* will make the results of research widely accessible and easy to understand, helping to realize a FAIR future for tissue imaging and digital pathology

### Box 1: Software for Managing and Visualizing Image Data

The OME-based *OMERO*^70^ server remains the most widely used image informatics system for microscopy data in a research setting; it is the foundation of the European Bioinformatics Institute (EBI)-based Image Data Resource (IDR)^33^, a prominent large-scale publicly accessible image repository, as well as more specialized repositories such as Pancreatlas^71^. *OMERO* has a client-server (three-tier) architecture involving a relational database, an image server, and one or more interoperable user interfaces. *OMERO* is well suited to managing image data and metadata and organizing images so that they can be queried using a visual index or via search^72^ (**Figure 2a**). In its current form, it does not perform sophisticated image analysis and no current browser is specialized for the creation of narrative guides.

A range of other software is available for static or partially interactive visualization of H&E and IHC images in a Web browser, including caMicroscope^73^ which is used to organize IHC and H&E images for The Cancer Genome Atlas (TCGA)^15^, The Cancer Imaging Archive (TCIA),^74^ Digital Slide Archive^37^, PathPresenter^75^, and Aperio^76^ (**Figure 2b**). Interested readers are referred to a recent White Paper from the Digital Pathology Association that discusses tools being developed to view bright-field digital pathology data^51^. However, such H&E and IHC viewers are not generally compatible with multi-channel images or to integrate different types of omics data.

### Box 2: Software for Analysis of Image Data

Multiple software suites have been developed for analysis of high-dimensional image data including *CellProfiler*^77^, *histoCAT*^78^, *Facetto*^79^, *QuPath*^80^, *Orbit*^81^, Mantis^82^, and *ASAP*^83^; many but not all run locally on the desktop (**Figure 2b**). These software systems generally perform image segmentation to identify individual cells or tissue-level features, determine cell centroids and shape parameters (e.g., area and eccentricity), and compute staining intensities in designated regions of an image and across all channels. The resulting vectors can then be processed using standard tools for high-dimensional data analysis such as supervised and unsupervised clustering, t-SNE^84^, or UMAP^85^ to identify cell types and study cell-cell interactions^86^. While some tools support segmentation, others require pre-generated segmentation masks and single-cell feature tables. As an alternative, emerging approaches analyze tissues at the level of individual pixels using machine learning and CNNs^87^. This approach potentially bypasses the need to segment individual cells from densely packed tissues in which cells can vary dramatically in size. A key feature of software such as *histoCat*^78^ or *Facetto*^88^ is integration of an image viewer and feature-based representations of the same data. This is essential for training and testing classifiers, quality-controlling image processing routines, and obtaining insight into spatial characteristics.

Commercial companies are also developing cloud-based computing platforms for digital pathology, including *HALO* (Indica Labs), *Visiopharm* (Visiopharm), and *PathViewer* (Glencoe Software, the commercial developer of *OMERO*). Academic (public domain) efforts include the *Allen Cell Explorer*^89^ and *napari*^90^ and build on highly successful open source software platforms such as *ImageJ*^91^. *Napari* will be particularly attractive to many computational biologists because it is based entirely in Python programming language and has both UI elements and a console. Commercial tools often strive for an all-in-one approach to analysis and visualization, but this comes at the cost of complexity, proprietary implementation, and licensing fees. It also ignores one of the primary lessons from genomics: progress in research rarely involves the use of a single integrated software suite, but instead an ecosystem of interoperable tools specialized to specific tasks.

## DATA AVAILABILITY

Source code for *Minerva Story* can be found at https://github.com/labsyspharm/minerva-story and detailed documentation and user guide at https://github.com/labsyspharm/minerva-story/wiki.

## ACKNOWLEDGEMENTS

This work was funded by NIH grants U54-CA225088 and U2C-CA233262 to P.K.S. and S.S., by U2C-CA233280 to P.K.S., and by the Ludwig Cancer Center at Harvard. The Dana-Farber/Harvard Cancer Center is supported in part by an NCI Cancer Center Support Grant P30-CA06516.

## OUTSIDE INTERESTS

PKS is a member of the SAB or BOD member of Applied Biomath, RareCyte Inc., and Glencoe Software, which distributes a commercial version of the OMERO database; PKS is also a member of the NanoString SAB. In the last five years the Sorger lab has received research funding from Novartis and Merck. Sorger declares that none of these relationships have influenced the content of this manuscript. SS is a consultant for RareCyte Inc. The other authors declare no outside interests.

